# The chromatin associated Sin3B is a critical regulator of DNA damage repair through the engagement of the Non-Homologous End Joining

**DOI:** 10.1101/2022.03.09.483624

**Authors:** Jorge Morales-Valencia, Alexander Calderon, Siddharth Saini, Gregory David

**Author notes:** Address correspondence to Gregory David, Ph. D.

## Abstract

Transcription and DNA damage repair act in a coordinated manner. The scaffolding protein Sin3B serves as a transcriptional corepressor of hundreds of cell-cycle-related genes. However, the role of Sin3B during the DNA damage response still remains unknown. Here, we show that Sin3B depletion delays the resolution of DNA double-strand breaks (DSBs) and sensitizes cells to diverse DNA-damaging agents, including cisplatin and doxorubicin. Furthermore, Sin3B is rapidly recruited to DNA damage sites and interacts with other DDR proteins. Mechanistically, Sin3B promotes non-homologous end joining DNA repair by directing the accumulation of MDC1 at DBSs. Altogether, our findings impute an unexpected function for the transcriptional co-repressor Sin3B as a gatekeeper of genomic integrity, and point to the inhibition of the Sin3B chromatin modifying complex as a novel therapeutic vulnerability in cancer cells.

## INTRODUCTION

Most cancer patients who develop advanced or metastatic disease are treated with combinations of radiotherapy (Brinkman, 2020) and conventional chemotherapeutic agents, such as anti-metabolites, alkylating agents (Fu, 2012) and topoisomerase inhibitors (Pommier, 2012). Such treatments lead to cytotoxicity in rapidly dividing cells by damaging the DNA and interfering with DNA transcription and cell division (Weber, 2014). However, increased tolerance to DNA damage in cancer cells combined with the genetic or epigenetic acquisition of resistance mechanisms can dramatically dampen the efficacy of these treatments. In particular, activation of DNA Damage Response (DDR) pathways promotes genotoxic resistance, which remains a major obstacle in successful cancer treatment (O’Connor, 2015). Thus, unveiling the mechanisms underlying DDR in cancer cells is paramount to improving the long-term efficacy of anti-tumor treatment regimens.

DNA damaging agents used for cancer treatment often induce double-strand breaks (DSBs), which are highly toxic lesions (Chapman, 2013). Despite their widely recognized cytotoxic effects, DSBs occur in physiological conditions, including during meiosis to generate genetic diversity by promoting exchange between homologous chromosomes (Youds, 2011) and in immune cells for the production of an immune repertoire during V(D)J recombination and class switch recombination (CSR) (Stavnezer, 2008). Thus, safeguard mechanisms are in place for DSBs to get resolved. In eukaryotes, the major repair mechanisms of DSBs consist of the homologous recombination (HR) pathway and the non-homologous end joining (NHEJ) pathway. HR accurately restores a damaged DNA sequence using the sister chromatid as a template and is therefore limited to S and G_2_ cell-cycle phases (Warmerdam, 2010). In contrast, NHEJ corrects DNA damage by directly ligating the two exposed DNA ends together, and is thus error-prone (Lord, 2012). NHEJ functions during all phases of the cell cycle and is the primary pathway of DSB repair in G_1_ phase cells (Huertas, 2010).

Effective DSBs repair depends on checkpoint mediators that coordinate cell cycle progression with DNA repair (Nyberg, 2002). Among these, 53BP1 plays a critical role in supporting effective NHEJ (Lukas, 2004). Upon DNA damage, ATM phosphorylates 53BP1 which then localizes to DSB sites to mediate recruitment of other repair factors (Anderson, 2001). Moreover, 53BP1 prevents the resection of DSBs in G_1_ phase of the cell cycle, and thus favors NHEJ over HR (Panier, 2014). MDC1 (Mediator of DNA Damage Checkpoint 1) acts as an upstream determinant of the interaction between 53BP1 and DSBs (Bekker-Jensen, 2005). Moreover, it has been suggested that MDC1 mediates NHEJ (Dimitrova, 2006), regulation of the DNA replication point (Wang, 2011) and mitosis (Townsend, 2009).

Multiple proteins bridging chromatin modifications and the DNA damage response contribute to the DSB recruitment and retention of additional factors contributing to the engagement of a specific repair program (Soria, 2012; van Attikum, 2009). Histone deacetylases HDAC1 and HDAC2 have been reported to be recruited to DNA damage sites and serve as important components of the DDR by regulating DSB repair, mainly by promoting effective NHEJ (Miller, 2010). Administration of HDAC inhibitors sensitizes cancer cells to DNA damaging therapies by altering chromatin structure and down-regulating DNA repair mechanisms (Kachhap, 2010; Robert, 2016). Consistently, several pharmacological inhibitors of HDACs are undergoing clinical testing for treatment of different types of cancer (McClure, 2018), but their efficacy remains limited due to the involvement of HDAC-containing complexes in virtually all chromatin based processes. HDACs are usually found as components of large multiproteic complexes that lack sequence-specific DNA-recognition domains. A major HDAC1/2 containing complex is the evolutionary conserved Sin3 complex (Silverstein, 2004). Mammalian cells express two Sin3 paralogs, Sin3A and Sin3B, which are recruited to target loci trough their association with sequence-specific transcription factors (van Oevelen, 2008; Bartke, 2010; Malovannaya, 2011). Our understanding of the biological functions of Sin3B remains incomplete, but recent evidence has implicated Sin3B in quiescence (Cantor, 2017; David, 2008), senescence (Grandinetti, 2009; Rielland; 2014; Bainor, 2017) and stem cell differentiation (Cantor, 2017). However, the contribution of Sin3B in the DDR in mammalian cells remained unexplored.

We report here that Sin3B-deleted cells are more sensitive to genotoxic stress, which correlates with an impaired recruitment of the DDR protein MDC1 and NHEJ-associated defects. Collectively, this data suggest that Sin3B could represent a target for adjuvant therapy to enhance anti-cancer chemotherapy response.

## RESULTS

### Sin3B protects against DNA damage-induced apoptosis

It had been previously reported that Sin3p/Rpd3p complex inactivation impairs survival in response to DNA DSBs in *S. cerevisiae* (Jazayeri, 2004). This observation led us to examine the chemo-sensitizing effect of Sin3B deletion in mammalian cells, by treating wild-type glioblastoma (T98G) cells and their Sin3B knock-out (Sin3B^-/-^) counterparts with escalating doses of cisplatin. Sensitivity to cisplatin was significantly increased in the absence of Sin3B, as evidenced by a 3-fold reduction in the IC50 (Fig 1A). This impaired cellular resistance to cisplatin treatment correlated with increased apoptosis in Sin3B^-/-^ cells compared to their wildtype counterparts (Fig 1B). Importantly, ectopic expression of Sin3B restored resistance to low doses of cisplatin treatment in Sin3B^-/-^ cells (Fig 1C) and reduced apoptosis levels upon treatment (Fig 1D), confirming that Sin3B inactivation is responsible for the abovementioned phenotypes. Cisplatin-induced apoptosis was detected in Sin3B^-/-^ cells as early as 12 hours following drug exposure and continued increasing over time, whereas wildtype cells did not exhibit any significant increase in cell death even after 40 hours of the same dose drug treatment (Fig S1A, B). Sensitization to cisplatin upon Sin3B inactivation was also confirmed in prostate (PC3) and breast (MCF7) cancer cells (Fig 1E, F). We additionally documented the enhanced sensitivity of Sin3B^-/-^ cells to the commonly used chemotherapy agent, doxorubicin, a topoisomerase inhibitor and DSBs inducer (Fig 1G-I) and to UV light (Fig S1C). Sin3B is a large scaffold protein devoid of enzymatic activity, whose three PAH domains, located in the N-terminal region, mediate protein-protein interactions. Mutation of the PAH1 or the PAH3 significantly altered the ability of exogenous Sin3B to reduce cisplatin-induced apoptosis in Sin3B^-/-^ cells, while PAH2 appeared dispensable (Fig S2A, B). Genetic inactivation of canonical Sin3-HDAC and DREAM complexes members, SAP30 and Lin9, respectively, was insufficient to sensitize cells to cisplatin (Fig S2C-F). Together, these results indicate that Sin3B protects from drug-induced cytotoxicity, independently of its ability to interact with its cognate partners.

**Figure 1.**
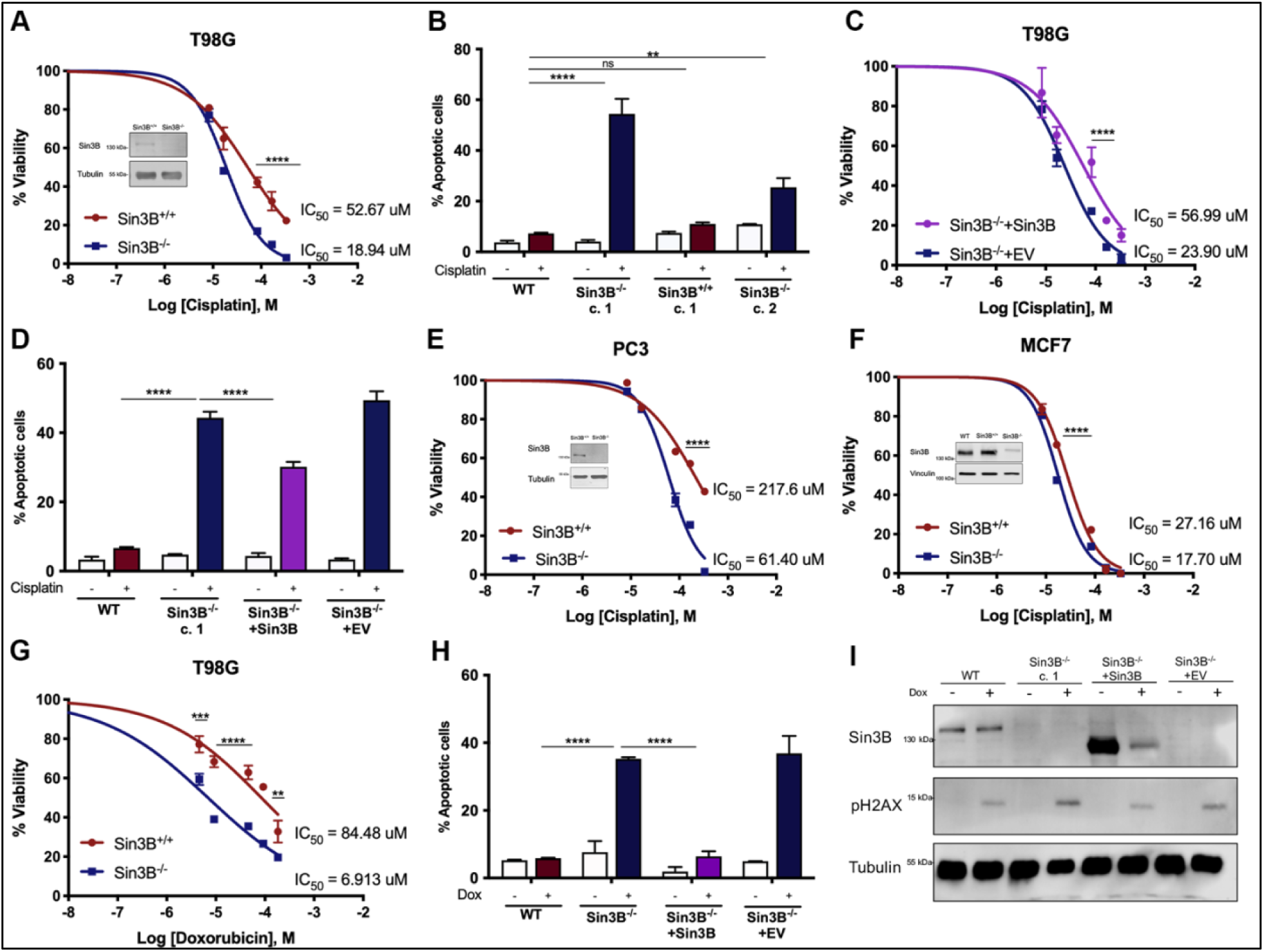
Sin3B protects against DNA damage-induced apoptosis. A. Sin3B^+/+^ and Sin3B^-/-^ T98G cells were treated with various concentrations of cisplatin for 12 h, then shifted to fresh medium and allowed to grow for 7 days before staining. Error bars represent SD; n = 3. Significance was determined by two-way ANOVA followed by a Sidak’s test. \*\*\*\**P*<0.0001. B. Sin3B^+/+^ and Sin3B^-/-^ T98G cells were treated with 25 ug/mL of cisplatin for 12 h. Cells were then subjected to Annexin V staining followed by flow cytometry analysis. Annexin V positive cells were then quantified. Error bars represent SD; n = 3. Significance was determined by two-way ANOVA followed by a Tukey’s test. \*\**P*<0.01, \*\*\*\**P*<0.0001. C. A T98G cell line stably expressing Flag-tagged Sin3B was generated. The resulting cells were treated with various concentrations of cisplatin for 12 h, then shifted to fresh medium and allowed to grow for 7 days before staining. Significance was determined by two-way ANOVA followed by a Sidak’s test. \*\*\*\**P*<0.0001. D. T98G cells expressing exogenous Sin3B were treated with 25 ug/mL of cisplatin for 12 h. Cells were then subjected to Annexin V staining followed by flow cytometry analysis. Annexin V positive cells were then quantified. Error bars represent SD; n = 3. Significance was determined by two-way ANOVA followed by a Tukey’s test. \*\*\*\**P*<0.0001. E. Sin3B^+/+^ and Sin3B^-/-^ T98G cells were treated with various concentrations of cisplatin for 12 h, then shifted to fresh medium and allowed to grow for 7 days before staining. Error bars represent SD; n = 3. Significance was determined by two-way ANOVA followed by a Sidak’s test. \*\*\*\**P*<0.0001. F. Sin3B^+/+^ and Sin3B^-/-^ MCF7 cells were treated with various concentrations of cisplatin for 12 h, then shifted to fresh medium and allowed to grow for 7 days before staining. Error bars represent SD; n = 3. Significance was determined by two-way ANOVA followed by a Sidak’s test. \*\*\*\**P*<0.0001. G. Sin3B^+/+^ and Sin3B^-/-^ T98G cells were treated with various concentrations of doxorubicin for 12 h, then shifted to fresh medium and allowed to grow for 7 days before staining. Error bars represent SD; n = 3. Significance was determined by two-way ANOVA followed by a Sidak’s test. \**P*<0.05, \*\**P*<0.01, \*\*\*\**P*<0.0001. H. Sin3B^+/+^ and Sin3B^-/-^ T98G cells were treated with 15 ug/mL of doxorubicin for 12 h. Cells were then subjected to Annexin V staining followed by flow cytometry analysis. Annexin V positive cells were then quantified. Error bars represent SD; n = 3. Significance was determined by two-way ANOVA followed by a Tukey’s test. \*\*\*\**P*<0.0001. I. Cells were treated with 15 ug/mL of doxorubicin for 12 h before being harvested for immunoblotting with the indicated antibodies.

### Sin3B-depleted cells exhibit defective DNA damage response

To gain insight into the contribution of Sin3B in the DNA damage response (DDR), we monitored DDR signaling in cisplatin-treated cells by documenting the emergence of yH2AX foci, a well-establish marker of DNA damage. Sin3B inactivation correlated with a nominal increase in yH2AX foci emergence in normal growing conditions. Strikingly, upon cisplatin exposure, Sin3B inactivated cells exhibited a significant increase in the number of yH2AX foci compared to their cisplatin-treated wildtype counterparts. Interestingly, the decrease in yH2AX foci number observed in wildtype cells during recovery from cisplatin exposure was also dampened in Sin3B^-/-^ cells (Fig 2A, B), suggesting that the presence of Sin3B may promote efficient repair of DNA breaks. Increased levels of yH2AX in Sin3B-null cells upon DNA damage was validated by Western Blot (Fig 2E). To directly assess the impact of Sin3B loss on DNA damage following cisplatin exposure, we performed comet assay analysis, which provides a qualitative assessment of the damaged genomic DNA. This analysis confirmed a significant increase in DNA damage levels in cells lacking Sin3B compared to control cells (Fig 2C, D). Furthermore, phosphorylation of the DNA damage sensing Chk2 was higher and more sustained in Sin3B^-/-^ cells compared to their wildtype counterparts following exposure to cisplatin (Fig 2E). Collectively, the above data indicates that Sin3B enables proper DSB repair.

**Figure 2.**
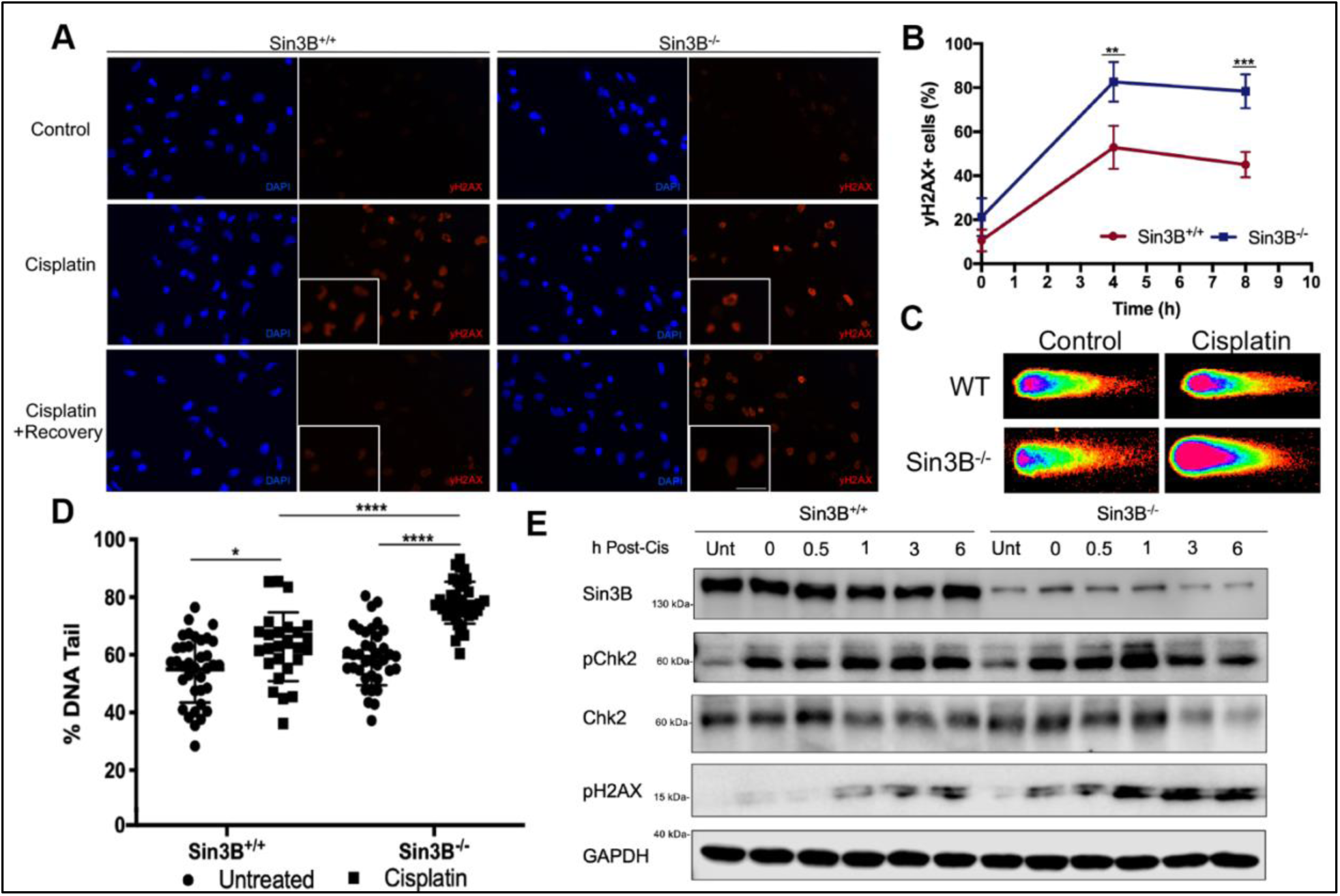
Sin3B-depleted cells exhibit defective DNA damage response. A. Cells were treated with 25 ug/mL of cisplatin for 4 h. Cells were fixed or given a recovery window of 4 hours where the drug was washed off and fresh media was added. Cells were then fixed, stained for yH2AX and imaged. Representative images of the data quantified in 2B. B. Quantification of yH2AX positive cells. Error bars represent SD; n = 3. Significance was determined by two-way ANOVA followed by a Sidak’s test. \*\**P*<0.01, \*\*\**P*<0.001. C. Representative images of tail moment from single-cell gel electrophoresis (neutral comet assay) of T98G cells treated with 25 ug/mL of cisplatin for 4 h. Drug was then washed off and fresh media was added for 4 additional hours. D. Tail moment quantification from single-cell gel of T98G cells. More than 200 total cells were considered. Error bars represent SD; n = 3. Significance was determined by two-way ANOVA followed by a Tukey’s test. \**P*<0.05, \*\*\*\**P*<0.0001. E. Cells were treated with 25 ug/mL of cisplatin for 4 h. Cells harvested for immunoblotting at different timepoints after drug removal.

### Sin3B is recruited to DNA Damage sites

To further elucidate the molecular bases for the contribution of Sin3B in DSB repair, we used live-cell imaging to track Sin3B localization as it relates to sites of DNA damage. Prior to the induction of DNA damage, green-fluorescent protein (GFP)-tagged Sin3B (Sin3B-GFP) exhibited a nuclear diffuse localization, consistent with what we and other have observed for endogenous Sin3B (data not show and Garcia-Sanz, 2014). However, Sin3B-GFP was rapidly recruited to sites of laser-induced DNA damage (Fig 3A, B). The molecular bases for the recruitment of Sin3B at DSB sites are elusive, prompting us to seek proteins that associate with Sin3B at DNA damage sites by a proximity-dependent biotin identification assay (BioID). We stably expressed biotin ligase-tagged Sin3B and confirmed its expression in cells treated with or without cisplatin (Fig 3C). Candidate protein interactors were isolated using streptavidin-conjugated beads and identified by mass spectrometry (MS). Consistent with previous reports, we identified canonical components of the mammalian homologs of the Rpd3 large and small complexes, as well as several components of the DREAM complex. Importantly, Sin3B-proximal proteins also include several proteins involved in the DDR, such as 53BP1, RIF1, MDC1, BRCA1/2 and others (Fig 3D). Gene Ontology analysis indicated that Sin3B-associated proteins identified by BioID, are enriched in the DDR pathways: Non-Homologous End Joining (NHEJ) and Homologous Recombination (HR) (Fig 3E). These results suggest that Sin3B is recruited to DNA damage sites and may interact with known DDR proteins.

**Figure 3.**
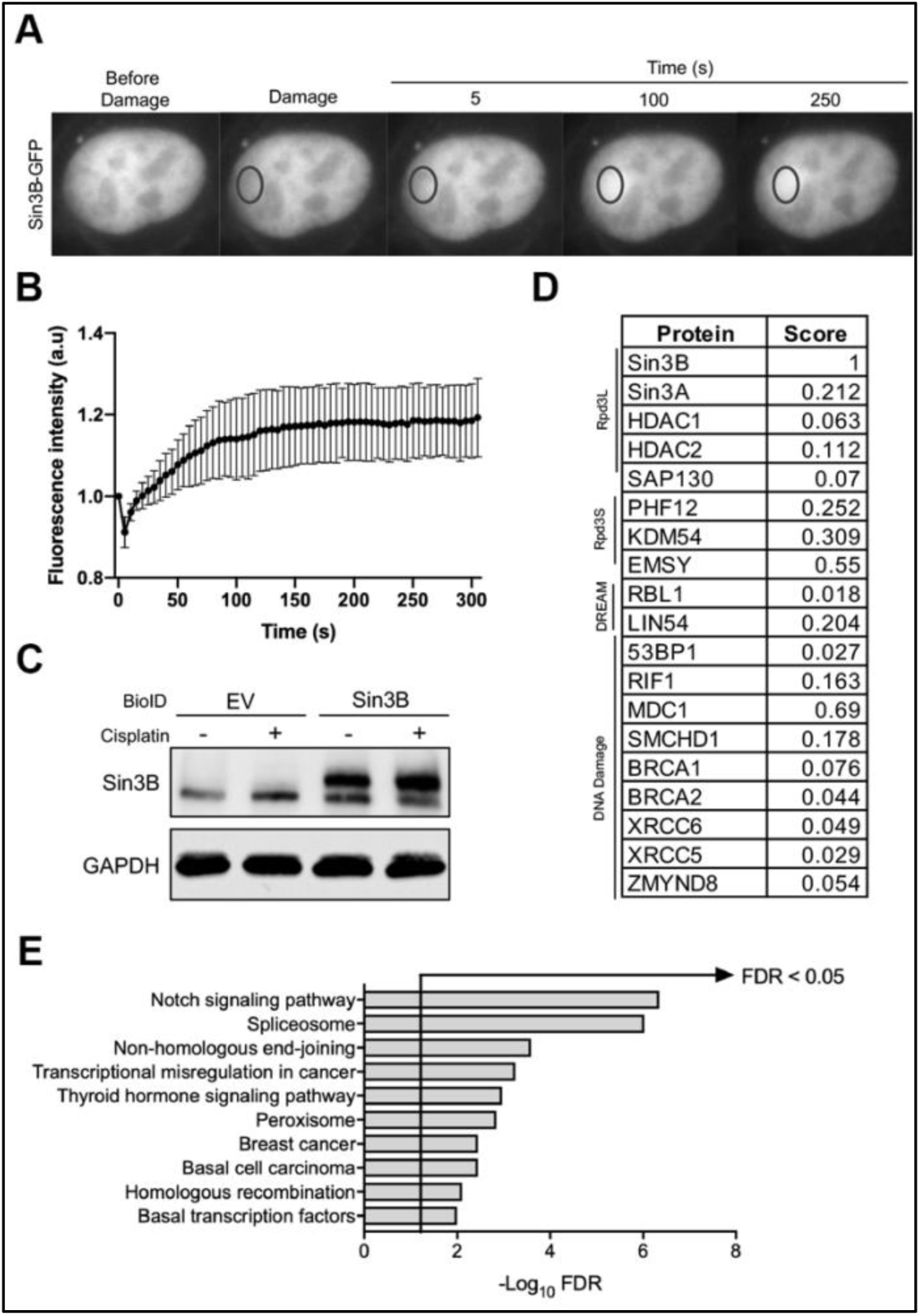
Sin3B is recruited to DNA Damage sites. A. Representative time-lapse images showing the accumulation of Sin3B-GFP to sites of laser-microirradiation (black circle). B. The difference in average fluorescence intensity in the damaged versus an undamaged region is plotted in relationship to time. A minimum of 50 different cells were analyzed. Error bars indicate SEM. C. Western blot shows stably expression of HA-Sin3B-BirA* in T98G cells in absence or presence of cisplatin (2.5 ug/mL for 12 h). D. Normalized score values of Sin3B interactors. The score is normalized to Sin3B signal. E. Gene ontology analysis of genes that were identified by BioID. Annotations were determined using Enrichr. Results with an FDR < 0.05 are shown.

### Sin3B facilitates the accumulation of MDC1 at DSBs

Mediator of DNA Damage Checkpoint 1 (MDC1) is tethered to sites of DNA damage and contributes to the recruitment of several other DDR proteins, including BRCA1, 53BP1 and the MRN complex (Stewart, 2003; Xu, 2003). Moreover, downregulation of MDC1 has been associated with hypersensitivity of cells to DSBs and aberrant apoptosis activation, reminiscent of the phenotypes elicited upon Sin3B inactivation (Coster, 2010). In light of the association we detected between Sin3B and MDC1 using the BioID approach, we sought to investigate the relationship between Sin3B and MDC1 recruitment at sites of DNA damage. First, we confirmed the interaction between ectopically expressed Sin3B and both endogenous and exogenous MDC1 by FLAG co-Immunoprecipitation (co-IP) (Fig 4A). Because MDC1 and H2AX extensively co-localize and display similar kinetics of foci formation upon DNA damage (Stewart, 2003), we examined the ability of MDC1 to form foci and colocalize with yH2AX in the absence of Sin3B. Sin3B-depleted cells exhibited a marked reduction in the colocalization of these two proteins upon cisplatin treatment (Fig 4B). Strikingly, Sin3B inactivation impaired MDC1 accumulation at DNA damage sites, as measured by live cell microscopy in laser micro irradiated MDC1-GFP cells (Fig 4C). Of note, MDC1 overexpression was not sufficient to rescue cisplatin sensitivity in Sin3B^-/-^ cells, suggesting that MDC1 requires Sin3B to promote DNA repair, or that Sin3B is required for the recruitment of additional DDR proteins, independently of MDC1 (Fig 4D). A known function of MDC1 in the response to DNA damage is its requirement for the intra-S phase checkpoint activation (Goldberg, 2003). Accordingly, while Sin3B^+/+^ cells accumulated in S phase when challenged with cisplatin, Sin3B^-/-^ cells presented with an intra-S phase checkpoint defect and accumulated in G_0_/G_1_ instead, similar to MDC1-null cells (Fig 4E and Goldberg, 2003). Collectively, these results point to an essential role for Sin3B in the recruitment of MDC1 to sites of DNA damage and the subsequent DDR.

**Figure 4.**
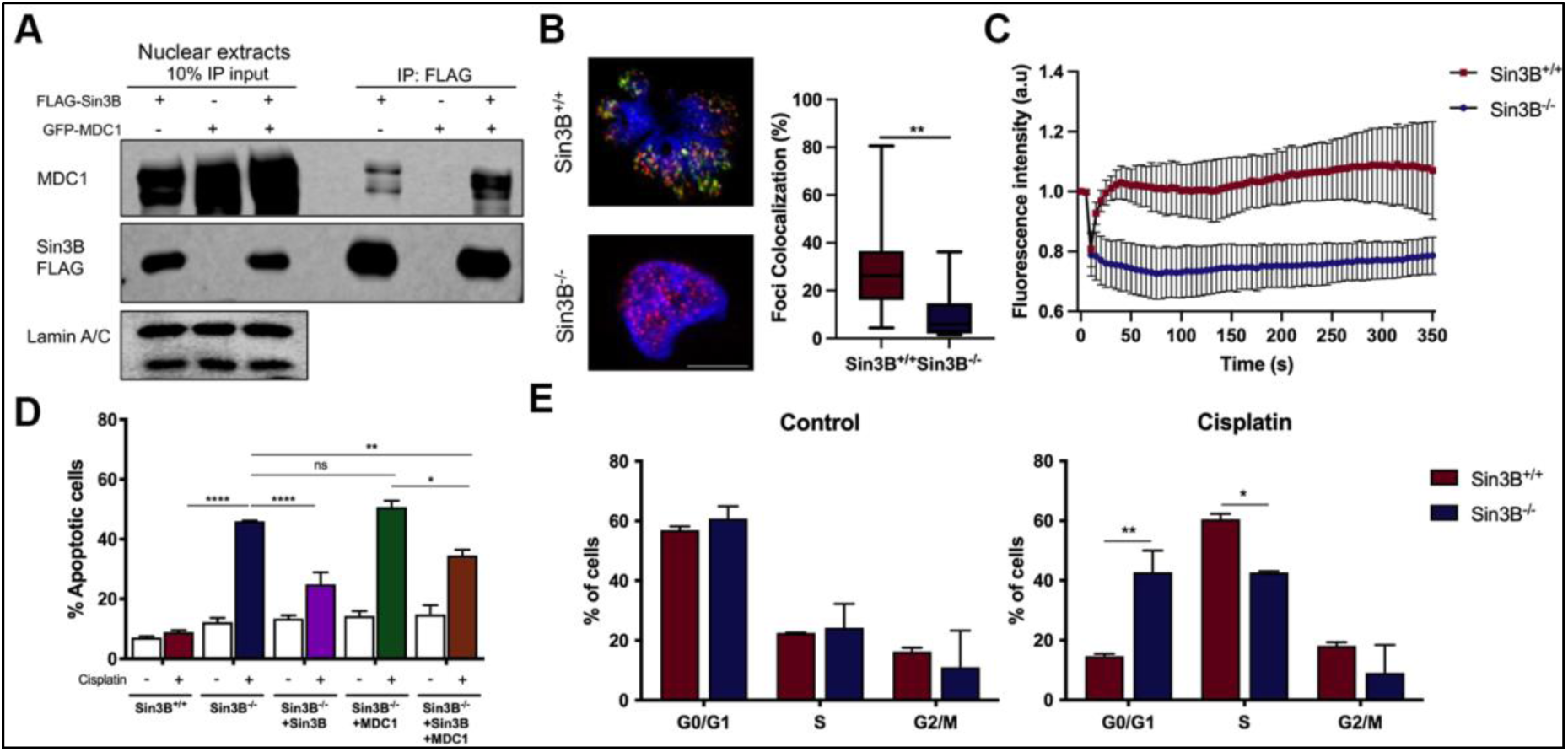
Sin3B facilitates the accumulation of MDC1 at DSBs. A. Co-immunoprecipitation with and anti-FLAG antibody of T98G cells transfected with MDC1-GFP. Immunoblots were performed using the indicated antibodies. 10% input. IP, immunoprecipitation. B. Sin3B^+/+^ and Sin3B^-/-^ T98G cells, expressing MDC1-GFP were treated with 25 ug/mL of cisplatin for 12 hours before being subjected to Annexin V staining followed by flow cytometry analysis. Annexin V positive cells were then quantified. Error bars represent SD; n = 3. Significance was determined by two-way ANOVA followed by a Tukey’s test. \**P*<0.05, \*\**P*<0.01, \*\*\*\**P*<0.0001. C. Accumulation of MDC1-GFP to sites of laser micro-irradiation. Difference in average fluorescence intensity in the damaged versus an undamaged region is plotted in relationship to time. A minimum of 50 different cells were analyzed. Error bars indicate SEM. D. Colocalization of MDC1/yH2AX in response to DNA damage. MDC1-GFP T99G cells were treated with 25 ug/mL of cisplatin for 4 h before being fixed, stained for yH2AX and imaged. Error bars represent SD; n = 3. Significance was determined by Student’s t-test. \*\**P*<0.01. E. Cell cycle distribution of Sin3B^+/+^ and Sin3B^-/-^ T98G cells as measured by PI incorporation after cisplatin treatment (2.5 ug/mL for 4 h). Error bars represent SD; n = 3. Significance was determined by two-way ANOVA followed by a Sidak’s test. \**P*<0.05, \*\**P*<0.01.

### cNHEJ is impaired upon Sin3B loss

Due to the known contribution of Sin3B on cell cycle progression, we sought to determine whether its chemo-protective properties coincided with the cells’ presence in a specific cell cycle phase. Using phase-specific blocking agents, we observed that Sin3B^-/-^ cells hypersensitivity to cisplatin was only evidenced in G_1_ phase (Fig 5A). This result raised the possibility that Sin3B protects cells from exogenous damage by modulating a DDR pathway active during the G_1_ phase of the cell cycle. Moreover, cell cycle exit induced by serum starvation conferred Sin3B^-/-^ cells a chemo-protective effect (Fig S3A). Canonical Non-Homologous End Joining (c-NHEJ) is responsible for DSB repair during G_1_ (Hustedt, 2016). 53BP1 is a critical factor for NHEJ (Gupta, 2014) and its accumulation at sites of DNA damage is dependent on MDC1 (Stewart, 2003; Eliezer, 2009). Thus, we aimed to determine the impact of Sin3B loss in the recruitment of 53BP1. Upon cisplatin exposure, we observed that 53BP1 foci number was decreased in Sin3B^-/-^ cells, compared to wildtype cells (Fig 5B, C), suggesting that Sin3B contributes to the recruitment of 53BP1 at DSBs. This result is consistent with the observation that, in addition to MDC1, Sin3B associates with 53BP1 (Fig 3D). NHEJ factors such as MDC1, RIF1 and 53P1 promote NHEJ and CSR (Lou, 2006; Manis, 2004; Ward, 2004). Therefore, we investigated the status of CSR in Sin3B^-/-^ mice by comparing the levels of serum immunoglobulins (Igs) to those of Sin3B^+/+^ animals. Following treatment, surface expression of IgG1 and IgG2b was reduced in Sin3B^-/-^ cultured B cells (Fig S4A, B). This reduction did not reach statistical significance. However, a similar defect in Ig class switching is observed for MDC1^-/-^ cells (Lou, 2006). To directly test the impact of Sin3B on c-NHEJ based repair of DSBs, we employed an I-Sce1 based c-NHEJ assay. Strikingly, genetic inactivation of Sin3B resulted in a decrease of GFP^+^ cells after nuclease induction, indicative of a defective c-NHEJ-based repair (Fig 5D). By contrast, Sin3B deletion did not prevent HR or altNHEJ, but instead resulted in an increased number of HR and alt-NHEJ events (Fig 5 E, F). This observation is consistent with published reports indicating that cancer cells become more reliant on specific DDR pathways when others are mutated or nonfunctional, as a compensatory response (Warmerdam, 2020). Finally, pharmacological inhibition of DNA-PK, a protein essential for effective c-NHEJ, synergized with cisplatin administration to induce apoptosis in wildtype cells. By contrast, these concomitant treatments merely resulted in an additive effect on apoptosis in Sin3B^-/-^ cells (Fig 5G). Taken together, these data indicate that Sin3B enables proper DDR signaling and repair by allowing efficient cNHEJ by promoting MDC1 accumulation and DSB sites.

**Figure 5.**
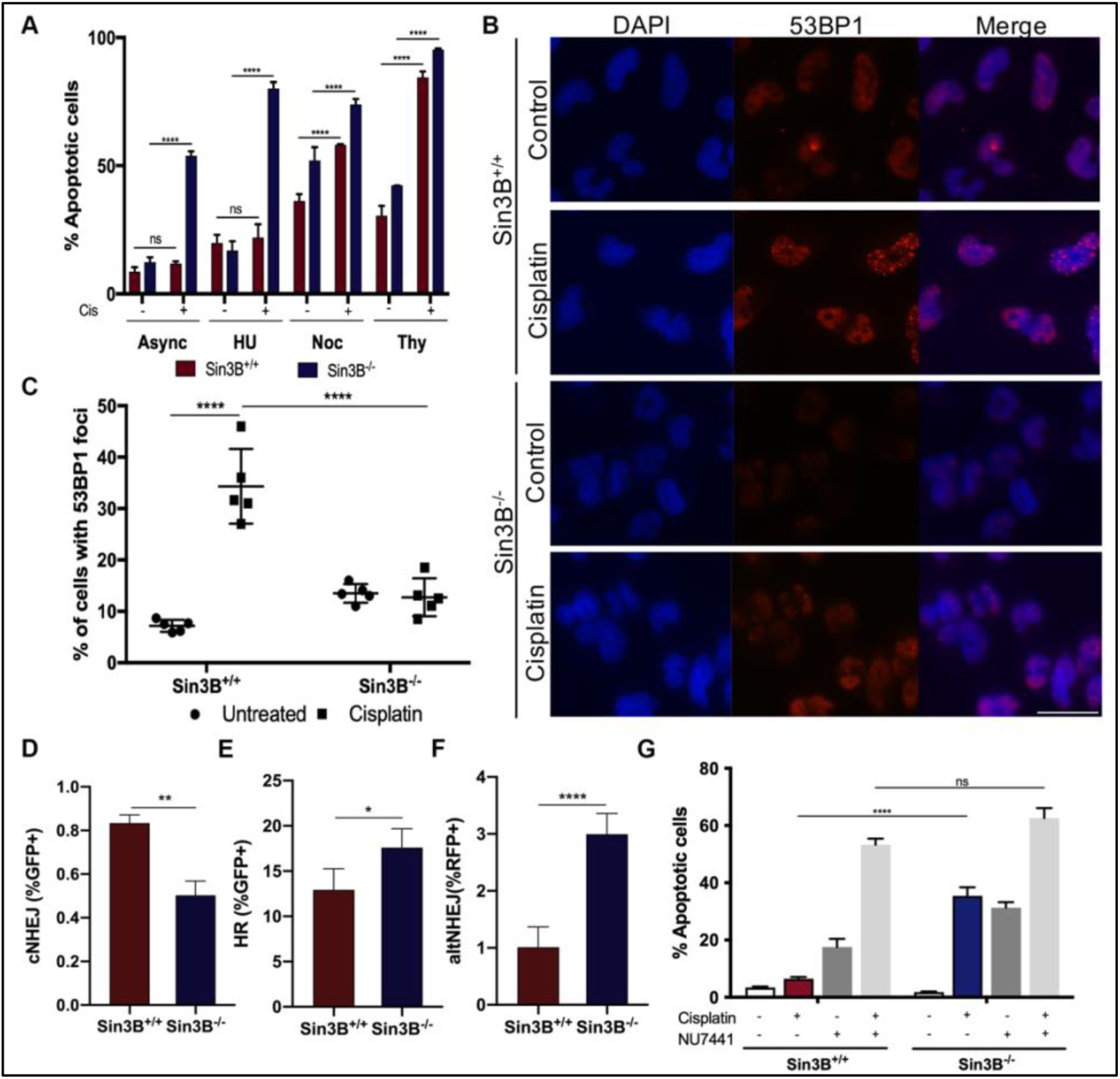
cNHEJ is impaired upon Sin3B loss. A. Sin3B^+/+^ and Sin3B^-/-^ T98G cells were synchronized in different phases of the cell cycle before being treated with 25 ug/mL of cisplatin for 12 followed by Annexin V staining followed by flow cytometry analysis. Annexin V positive cells were then quantified. Error bars represent SD; n = 3. Significance was determined by two-way ANOVA followed by a Tukey’s test. \*\*\*\**P*<0.0001. B. Cells were treated with 25 ug/mL of cisplatin for 4 h. Cells were then fixed, stained for 53BP1 and imaged. Representative images of the data quantified in 2C. C. Quantification of cells with 53BP1 foci. Error bars represent SD; n = 3. Significance was determined by two-way ANOVA followed by a Sidak’s test. \*\*\**P*<0.001. D. T98G cells containing the EJ5-GFP reporter were analyzed for GFP positivity, 72 hours after nuclease induction. The percentage of GFP^+^ cells were determined for each individual condition and subsequently normalized to transfection efficiency. Error bars represent SD; n = 3. Significance was determined by Student’s t-test. \*\**P*<0.01. E. T98G Sin3B^+/+^ and Sin3B^-/-^ TLR cell lines were created and then infected with lentivirus containing SceI and the template for eGFP reconstitution. After 72 hours, percentage of GFP^+^ cells were determined by flow cytometry for each individual condition and subsequently normalized to infection efficiency. Error bars represent SD; n = 3. Significance was determined by Student’s t-test. \**P*<0.05. F. Similar to E, RFP^+^ cells were determined by flow cytometry. Error bars represent SD; n = 3. Significance was determined by Student’s t-test. \*\*\*\**P*<0.0001. G. Sin3B^+/+^ and Sin3B^-/-^ T98G cells were treated with 25 ug/mL of cisplatin for 12 h and 10 nM NU7441. Cells were then subjected to Annexin V staining followed by flow cytometry analysis. Annexin V positive cells were then quantified. Error bars represent SD; n = 3. Significance was determined by two-way ANOVA followed by a Tukey’s test. \*\*\*\**P*<0.0001.

## DISCUSSION

Altogether, our data point to a model where Sin3B regulates DNA damage repair through cNHEJ. The inefficient repair resulting from the loss of Sin3B causes the accumulation of DNA damage and ultimately leading to cell death upon exposure to genotoxic drugs. These results resonate with previous observations made in budding yeast, where sin3 and rpd3 mutant strains display reduced viability upon exposure to the DBS-inducing drug, phleomycin (Jazayeri, 2004). Moreover, Sin3B is a downstream target of the Polycomb protein Bmi-1, which has been proposed to facilitate the repair of DSBs (Facchino, 2010; DiMauro, 2015; Lin, 2015). Additional reports indicate that human HDAC1 and HDAC2 function in the DDR to promote NHEJ (Miller, 2010), but the identity of the HDAC-containing complex(es) responsible for these effects is unknown.

While our experiments clearly indicate a role for Sin3B in facilitating NHEJ, the exact molecular mechanism by which Sin3B supports this effect remained elusive. In yeast, Sin3p is required for the deacetylation of H4K16 in the vicinity of a DSB believed to result in the formation of a localized hypoacetylated chromatin domain (Jazayeri, 2004). Such hypoacetylated domains have been also found at sites of DSBs in mammalian cells (Shanbhag, 2010), pointing to the possibility that Sin3B functions in DDR through the tethering of a H3K16 deacetylase activity at damaged loci. However, we observed that, while Sin3B inactivation resulted in higher H4K16 acetylation in normal growing conditions, induction of a DSB did not exacerbate this difference (Fig S5 A, B). This raised the possibility that Sin3B’s role in promoting cNHEJ might be independent of its ability to tether histone modifying activities at damaged loci. An alternative explanation for the decrease in NHEJ efficiency upon Sin3B deletion could be related to an impaired regulation of cell cycle progression. We and other have shown that loss of Sin3B primes cells for cell cycle entry through DREAM target genes de-repression (Rayman, 2012; Bainor, 2018). These observations raise the possibility that Sin3B^-/-^ cells progress through G_1_ too rapidly to ensure the proper engagement of the NHEJ pathway. However, genetic inactivation of Lin9, a conserved subunit of the DREAM complex, did not recapitulate the phenotype observed in Sin3B^-/-^ cells (Fig S2E, F). It is therefore unlikely that the sensitization to genotoxic drugs observed in Sin3B^-/-^ cells could be solely explained by the impact of Sin3B on cell cycle progression. Rather, our results point to a model where Sin3B modulates NHEJ by serving as a scaffold and coordinate the recruitment of multiple DDR-related proteins, including MDC1. Strikingly, Sin3B inactivation drastically alters the accumulation of MDC1 at DNA damage sites. MDC1 recruitment to DSBs is required for activation of the intra-S-phase DNA damage checkpoint necessary to protect genomic integrity and ensure replication fidelity (Goldberg, 2003). Similarly, we have observed that Sin3B enables S-phase arrest caused by cisplatin (Fig 4E). Interestingly, MDC1 inactivation has been reported to lead to sensitivity to cisplatin and doxorubicin in different cancer types including breast (Wang, 2018), cervical (Yuan, 2012), nasopharyngeal (Zeng, 2016) and esophageal (Yang, 2010). We believe Sin3B regulates DDR by tethering large complex(es) at sites of DSB, reminiscent but not overlapping with its function in transcriptional repression. Additional work is necessary to characterize said complexes and the DNA region that they bind as well as the chromatin modifications associated with them since Sin3B harbors the ability to interact with multiple chromatin modifiers.

As cancer cells overly rely on DNA repair pathways to overcome damage triggered by uncontrolled replication, the development of new treatment modalities targeting specific DNA repair pathways aimed at improving gold standard treatments represent a promising therapeutic opportunity. Several lines of evidence suggest that the targeting of core NHEJ factors offers a therapeutic window, making this pathway an ideal target (Marangoni, 2000; Sishc, 2017). The frequent emergence of resistance to anti-cancer targeted therapies warrants the characterization of unsuspected targets in well-delineated molecular pathways. Based on this study, we propose that Sin3B represents a target for adjuvant therapy to enhance anti-cancer chemotherapy response. Intriguingly, administration of Sin3-interacting domain (SID) decoys, which block the association between Sin3 proteins and specific interactors, can inhibit breast cancer growth in mouse models (Farias, 2010). Overall, an improved understanding of the composition and function of specific Sin3B-containing complexes should enable the generation of novel therapeutic strategies for the treatment of cancer.

## MATERIALS AND METHODS

### Cell culture, reagents and treatments

T98G and MCF7 cells were cultured in DMEM (Cellgro), 10% fetal bovine serum, and 1% penicillin/streptomycin (Cellgro). PC3 cells were cultured in Ham’s F-12K medium (Gibco), 10% fetal bovine serum and 1% penicillin/streptomycin (Cellgro). All cultures were maintained in 5% CO2 at 37°C. Unless indicated otherwise, cells were treated with 25 ug/mL cisplatin (Sigma) or 15 ug/mL doxorubicin (Sigma) overnight. For DNA-PK inhibition, cells were treated with 10 nM NU7441 (STEMCELL) for 30 minutes before being treated with cisplatin.

### Establishment of Sin3B null CRISPR cell lines

Cells were transfected with two independent sgRNA guide pairs, which targeted exon 2 of Sin3B, cloned into the lentiCRISPR v2 plasmid, gift from R. Possemato, NYU School of Medicine, New York, NY. Cells were selected (1 ug/ml of puromycin), clonally amplified and assessed for Sin3B deletion by Western Blot.

### Cell survival assays

2000 cells were plated in triplicate in 96-well plates and allowed to adhere overnight. Cells were then treated with increasing concentrations of cisplatin or doxorubicin overnight. Wells were washed with PBS and cells were fixed with 2% glutaraldehyde in PBS for 15 minutes. Cells were then stained with crystal violet (0.1% in 10% ethanol) for 30 minutes. After washing and drying, cells were distained in 10% acetic acid for 15 minutes. Optical density (OD) was measured at 595 nm absorbance.

### Annexin V-apoptosis assay

Treated and untreated cells were collected, without washing to collect all floating cells, centrifuged at 1300 rpm for 3 minutes. After discarding supernatant, 5 uL of Annexin V (BioLegend) was added to cells resuspended in 200 uL of binding buffer (BioLegend). Cells were then incubated for 30 minutes at room temperature in the dark before centrifuging them again. Cells were resuspended in 200 uL of binding buffer after discarding the supernatant and analyzed via flow cytometry using a FACSCalibur Flow Cytometer (BD).

### Protein extracts and Western blotting

Cells were lysed in 1× RIPA buffer (1% NP-40, 0.1% SDS, 50 mM Tris-HCl, pH 7.4, 150 mM NaCl, 0.5% sodium deoxycholate, 1 mM EDTA), 0.5 μM DTT, 25 mM NaF, 1 mM sodium vanadate, 1 mM PMSF, and cOmplete protease cocktail inhibitor (Sigma). Samples were resolved by SDS-PAGE and analyzed by standard western blotting techniques. The following primary antibodies were used: mouse anti-tubulin (Sigma T9026), mouse anti-vinculin (Sigma V9131), rabbit anti-Sin3B (Santa Cruz AK-12), rabbit anti-γ-H2AX (Cell Signaling 9718), mouse anti-GAPDH (Sigma MAB374), rabbit anti-SAP30 (Upstate 06-875), mouse anti-Lin9 (Santa Cruz sc-398234), rabbit anti-Chk2 (Cell Signaling 2662T), rabbit anti-pChk2 (Cell Signaling 2661T), rabbit anti-MDC1 (Abcam ab11171), mouse anti-Lamin A/C (Santa Cruz sc-376248).

### Immunofluorescence analyses

Cells were plated in duplicate on coverslips. Treated and untreated cells were fixed with 4% paraformaldehyde–PBS for 15 min and blocked with 0.3% Triton X-100–5% goat serum–PBS for 1 h. Cells were then incubated with rabbit anti-γ-H2AX (Cell Signaling 9718) at 1:400 dilution or rabbit anti-53BP1 (Cell Signaling 2675S) at 1:500 dilution in 0.3% Triton X-100–1% BSA–PBS overnight at 4°C. The following day, cells were incubated with 1:500 Alexa Fluor 488–anti-rabbit secondary antibody (Life Technologies) in 0.3% Triton X-100–1% BSA–PBS for 1 h. Coverslips were then washed and mounted as described above. Slides were examined on a Zeiss AxioImager A2 microscope.

### Comet assay

Assay was performed using the Single Cell Gel Electrophoresis kit (Trevigen). Treated and untreated cells were suspended at 1×10^5^ cells/ml in ice cold Ca^+2^ and Mg^+2^ free-PBS. Cells were combined with molten LMAgarose (at 37°C) at a ratio of 1:10 (v/v) and 50 µl were transferred onto a slide. Slides were incubated at 4°C in the dark for 10 minutes before being immersed in 4°C Lysis Solution for 1 hour. Slides were then placed in the electrophoresis tank and incubated with electrophoresis solution for 30 minutes before running electrophoresis for 30 minutes (25 V, 1.25 V/cm). Slides were dried before staining with SYBR®gold (Sigma). Comets were imaged using a Zeiss AxioImager A2 microscope and scored using the software CometScore 2.0.

### Laser micro-irradiation

Damage was created by exposure to a UV-A laser beam in cells plated on glass-bottomed dishes (Greiner Bio-One 543979). Cells were pre-sensitized with 1 ng of Hoescht in normal medium for 20 minutes at 37°C. Laser micro-irradiation was carried out with a FluoView 1000 confocal microscope (Olympus) and a 405 nm laser diode (6 mW). Laser settings (0.4 mW output) were used to generate DNA damage that was restricted to the laser path in a pre-sensitization-dependent manner. Variation of the fluorescence intensity was quantified as the difference between the average fluorescence intensity in the damaged region versus the average fluorescence intensity in an undamaged region of the same size in the same cells. Each curve represents data obtained from at least 10 independently filmed cells.

### BioID

Protein-Proximity labeling in living cells was done as previously described (Sears, 2020). Briefly, when cells expressing the construct were around 80% confluent, media was replaced and biotin was added to a final concentration of 50 uM for 16-18 hours. Cells were then collected, lysed and sonicated. A pre-clearing step was performed to remove proteins that were no specifically bound. To isolate biotinylated proteins, the cell lysate was incubated with streptavidin-conjugated sepharose beads. Beads were then washed 5 times and finally resuspended in 50 uL 50 mM ammonium bicarbonate. Isolated biotinylated proteins were resolved by SDS-PAGE and analyzed by silver staining, before sending samples for MS analysis.

### Nuclear lysate extraction and immunoprecipitation

Nuclear extracts were prepared from 2×10cm dishes of T98G cells stably expressing Sin3B-FLAG and/or MDC1-GFP. Nuclei were extracted by quick lysis in RSB-G40 lysis buffer (10mM Tris 7.4, 10mM NaCl, 3mM MgCl_2_, 10% glycerol, 0.25% NP-40 and supplemented with 0.5 mM DTT, 25 mM NaF, 1mM sodium orthovanadate, 1mM PMSF, and cOmplete protease inhibitor cocktail (Sigma). Nuclei were washed with RSB-G40 without NP-40, and lysed with high-salt extraction buffer (20 mM HEPES pH7.9, 420mM NaCl,1.5 mM MgCl_2_, 0.2 mM EDTA, 25% glycerol with additives. Salt concentrations were adjusted with the addition of No salt binding buffer (20mM HEPES pH7.9, 5 mM MgCl_2_, 10% glycerol, 0.1% Tween-20 with additives), and lysates were subjected to immunoprecipiation by incubation with EZview NRed ANTI-FLAG M2 affinity gel beads (Sigma) overnight. Immunoprecipitates were washed 3 times with binding buffer (20mM HEPES pH 7.9, 0.1 M NaCl, 5mM MgCl_2_, 10% glycerol, 0.1 % Tween-20 with additives) and FLAG beads were eluted by boiling in 1×SDS/Laemmli buffer (62.5 mM Tris HCl, pH 6.8, 2% SDS, 10% (v/v) glycerol, 0.002% bromophenol blue, 5% 2-Mercaptoethanol). Immunoprecipitates and 10% IP input lysates were loaded on 5.5% SDS-PAGE gel and probed with indicated antibodies: rabbit anti-Sin3B (Novus NBP2-20367), rabbit anti-MDC1 (Abcam ab11171), mouse anti-Lamin A/C (Santa Cruz sc-376248).

### Cell cycle analysis

Cells were harvested and washed with PBS before being fixed by transferring them drop wise into tubes containing 70% ethanol. Cells were stored at 4 C for at least 2 h. Samples were then centrifuged and washed with PBS twice before being resuspended in propidium iodide staining (10 ug/mL PI, 0.1% Triton X-100 and 50 ug/mL of DNAse-free RNAse A in PBS). Samples were incubated for 30 minutes at room temperature before being analyzed by flow cytometry.

### cNHEJ assay

pimEJ5GFP reporter system, which allows us to monitor the end joining between two distal tandem I-SceI recognition sites that restores GFP expression, was employed. T98G cells were co-transfected with 1ug/well of the pimEJ5GFP (Addgene #44026) and ISceI-GR-RFP (Addgene #17654) plasmids. The following day cells were treated with 0.2 mM TCA to induce activation of the nuclease. Seventy-two hours later, cells were harvested and analyzed for GFP positive cells by flow cytometry. For each analysis, 1×10^4^ cells were collected, and each experiment was repeated three times. Results were normalized to transfection efficiency.

### HR/altNHEJ assay

We employed the Traffic Light Reporter assay (Addgene #31482) to look at HR/altNEHJ efficiency. Briefly, a DSB is induced by SceI, if the break is resolved through the HR pathway, the full eGFP sequence will be reconstituted, and cells will fluoresce green; if the break undergoes altNHEJ, eGFP will be translated out of frame and the T2A and mCherry sequences are rendered in frame to produce red fluorescent cells. T98G Sin3B^+/+^ and Sin3B^-/-^ TLR cell lines were created and then infected with lentivirus containing SceI and the template for eGFP reconstitution (Addgene #32627), after 72 hours, flow cytometric analysis was performed.

### *Ex vivo* B cell class switch recombination

B cells were purified by negative selection from single-cell suspensions from spleen using magnetic separation B cell isolation kit (Miltenyi Biotec 130-090-862). Purified cells were cultured for 4 days in RPMI (Corning, 10-040-CV) supplemented with 10% FBS, LPS (20 ug/ml, Sigma L4130) and/or IL-4 (10 ng/ml, PreproTech #AF-214-14). Surface immunoglobulin expression was then assessed by flow cytometry.

### siRNA Transduction

Transient siRNA transfections were carried out with Lipofectamine RNAiMax (Invitrogen) according to manufacturer’s instruction. Analysis were performed 72 hours after siRNA transfection.

### H4K16 ChIP Qpcr

Chromatin immunoprecipitation (ChIP) was carried out as described previously (Shanbhag, 2010). Induction of DSBs in U2OS-DSB reporter cells line was induced by addition of Shield-1 (1 uM, TAKARA 632189) and 4-OHT (200 nM, Sigma 68392-35-8), this system uses an mCherry-LacI-FokI nuclease fusion protein to create DSBs within a single genomic locus (Tang, 2012). Cells were cross-linked with 1% (v/v) formaldehyde for 10 min, followed by the addition of glycine to 0.125 M for 5 min to stop the cross-linking. Cells were lysed in 10 mM Tris, pH 8.0, 10 mM NaCl, 0.2% NP-40. Nuclei were isolated, resuspended in 50 mM Tris pH 8.1, 10 mM EDTA, 1% SDS and sonicated to obtain approximately 200–500 bp chromatin fragments using a Bioruptor (Diagenode). Chromatin fragments were precleared with salmon sperm DNA (Trevigen, 9610-5-D)/protein-A agarose (Santa Cruz sc-2003) and incubated with H4K16Ac antibody (Millipore 07-329)-protein A agarose overnight at 4°C. Beads were washed once in low salt buffer (20 mM Tris, pH 8.1, 2 mM EDTA, 50 mM NaCl, 1% Triton X-100, 0.1% SDS), twice in high salt buffer (20 mM Tris, pH 8.1, 2 mM EDTA, 500 mM NaCl, 1% Triton X-100, 0.1% SDS), once in LiCl buffer (10 mM Tris, pH 8.1, 1 mM EDTA, 0.25 mM LiCl, 1% NP-40, 1% deoxycholic acid) and twice in TE buffer (10 mM Tris-HCl, pH 8. 0, 1 mM EDTA). Washed beads were eluted twice with 100 uL of elution buffer (1% SDS, 0.1 M NaHCO3) and de-crosslinked (0.1 mg/ml RNase, 0.3 M NaCl and 0.3 mg/ml Proteinase K) overnight at 65°C. The DNA samples were purified with Qiaquick PCR columns (Qiagen 20104). qPCR was carried out on a QuantStudio™ 5 instrument using the SYBR Green detection system. ChIP-qPCR primer sequences used are forward: GGAAGATGTCCCTTGTATCACCAT and reverse: TGGTTGTCAACAGAGTAGAAAGTGAA.

### Statistical analysis

Results were compared statistically using GraphPad Prism software. Values were subjected to unpaired two-tailed t tests, multiple t tests, one-way analysis of variance (ANOVA) followed by Dunnett’s multiple-comparison test, or two-way ANOVA followed by Tukey’s multiple-comparison test. Results were considered significant at p < 0.05 and are presented as mean ± SEM.

## ACKNOWLEDGEMENTS

The authors sincerely thank all members of the David lab for helpful discussions during the preparation of this manuscript. We thank the NYU Proteomics Laboratory for help with mass spectrometry analysis. We thank Dr. Michele Pagano (NYU School of Medicine) and Dr. Richard Possemato (NYU School of Medicine) and members of their labs for the generous gift or reagents and plasmids as well as Dr. Roger Greenberg for the U2OS-DSB reporter cells (University of Pennsylvania). Finally, we thank Dr. Susan Logan (NYU School of Medicine), Dr. Tony Huang (NYU School of medicine), Dr. Richard Possemato (NYU School of Medicine) and Dr. Eli Rothenberg (NYU School of medicine) for helpful discussions. This work was funded by NIH/NCI (CA246416) [GD], NYS DoH (C36617GG) [GD] and the NYSTEM Institutional Training Grant (C322560GG) [JMV].

## AUTHOR CONTRIBUTIONS

JM-V designed and performed the experiments presented in this manuscript and analyzed the data. AC performed confocal imaging and the *ex vivo* B-cells class switch recombination experiment. SS performed co-immunoprecipitation experiments. GD design the study and supervised the research. JM-V and GD wrote the manuscript.

## SUPPLEMENTAL FIGURES

**Figure S1.**
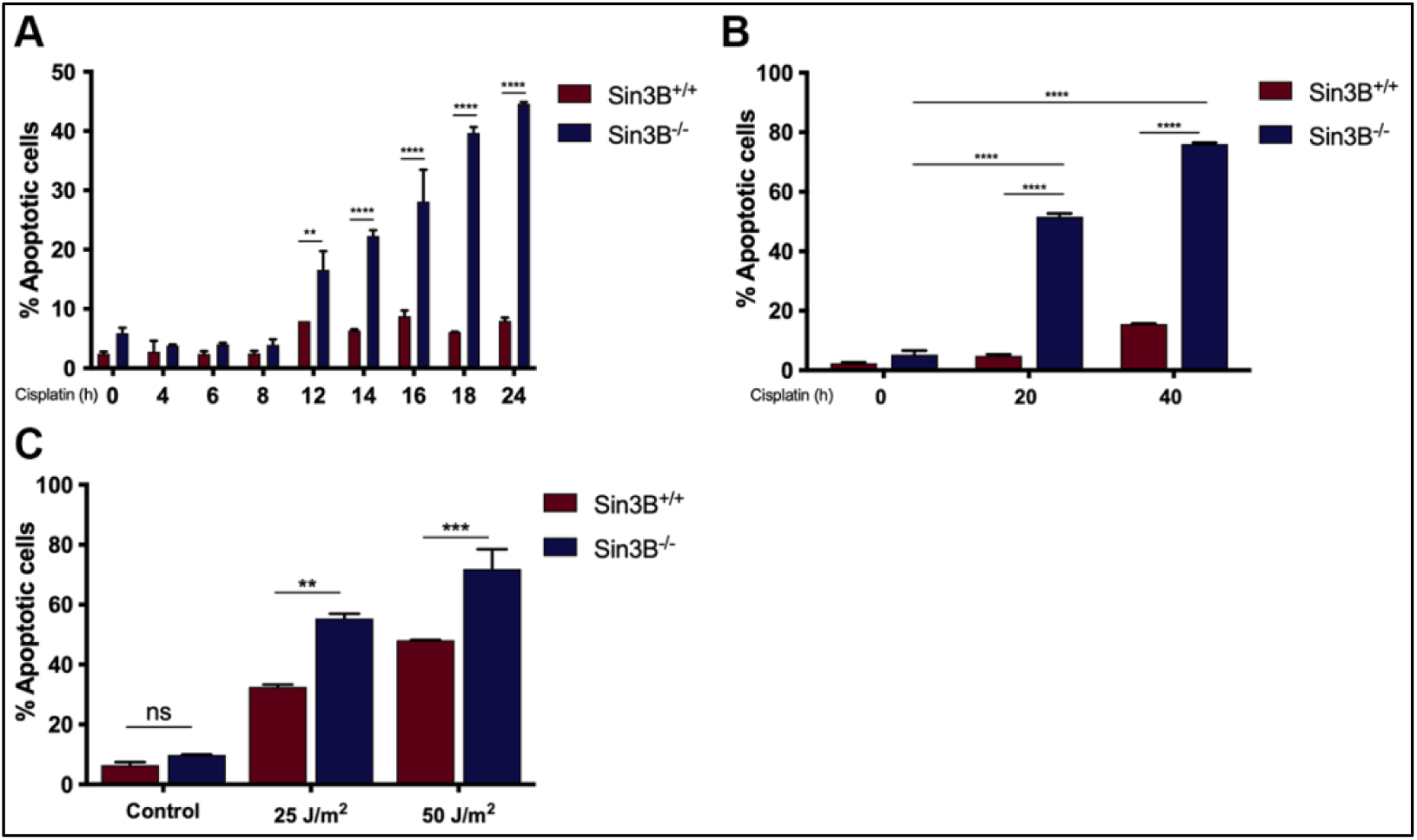
Sin3B protects against DNA damage-induced apoptosis. A. Sin3B^+/+^ and Sin3B^-/-^ T98G cells were treated with 25 ug/mL of cisplatin for different periods of time ranging from 0 to 24 hours. Cells were then subjected to Annexin V staining followed by flow cytometry analysis. Annexin V positive cells were then quantified. Error bars represent SD; n = 3. Significance was determined by two-way ANOVA followed by a Tukey’s test. \*\**P*<0.01, \*\*\*\**P*<0.0001. B. Sin3B^+/+^ and Sin3B^-/-^ T98G cells were treated with 25 ug/mL of cisplatin for 20 or 40 hours. Cells were then subjected to Annexin V staining followed by flow cytometry analysis. Annexin V positive cells were then quantified. Error bars represent SD; n = 3. Significance was determined by two-way ANOVA followed by a Tukey’s test. \*\*\*\**P*<0.0001. C. Sin3B^+/+^ and Sin3B^-/-^ T98G cells were exposed to UV light radiation. After 48 hours, cells were then subjected to Annexin V staining followed by flow cytometry analysis. Annexin V positive cells were then quantified. Error bars represent SD; n = 3. Significance was determined by two-way ANOVA followed by a Tukey’s test. \*\*\*\**P*<0.0001.

**Figure S2.**
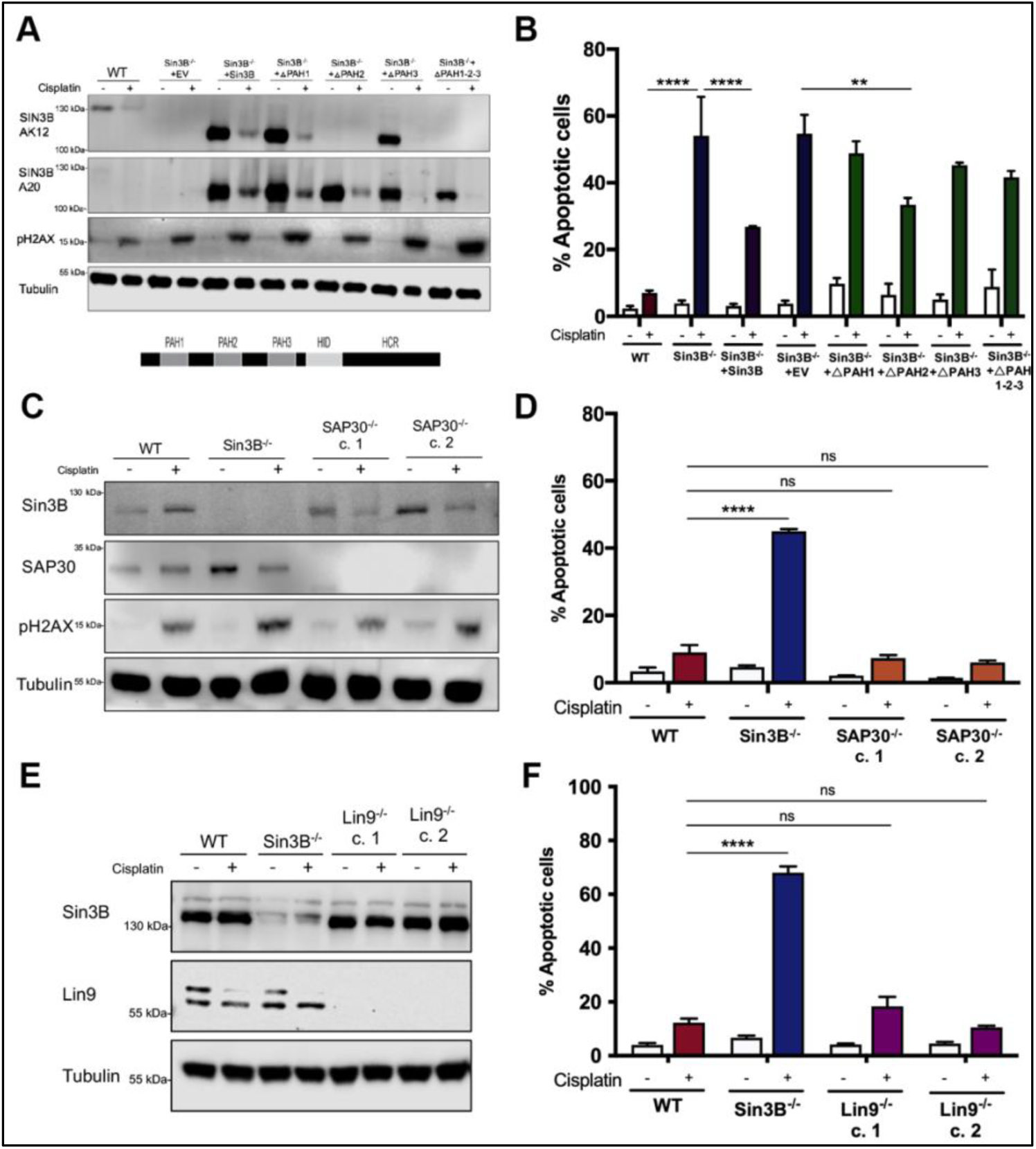
Sin3B protects from drug-induced cytotoxicity, independently of its ability to interact with its cognate partners. A. Western blot on whole-cell lysate from Sin3B^+/+^ and Sin3B^-/-^ T98G cells expressing different Sin3B truncation mutants probed using the indicated antibodies. Sin3B AK12 antibody targets the PAH2 domain of Sin3B and probing with Sin3B A20 was necessary to confirm expression. B. Sin3B^+/+^ and Sin3B^-/-^ expressing different Sin3B truncation mutants T98G cells were treated with 25 ug/mL of cisplatin for 12 hours. Cells were then subjected to Annexin V staining followed by flow cytometry analysis. Annexin V positive cells were then quantified. Error bars represent SD; n = 3. Significance was determined by two-way ANOVA followed by a Tukey’s test. \*\*\*\**P*<0.0001. C. Western blot on whole-cell lysate from Sin3B^+/+^, Sin3B^-/-^ and SAP30^-/-^ T98G cells expressing different Sin3B truncation mutants probed using the indicated antibodies. D. Sin3B^+/+^, Sin3B^-/-^ and SAP30^-/-^ T98G cells were treated with 25 ug/mL of cisplatin for 12 hours. Cells were then subjected to Annexin V staining followed by flow cytometry analysis. Annexin V positive cells were then quantified. Error bars represent SD; n = 3. Significance was determined by two-way ANOVA followed by a Tukey’s test. \*\*\*\**P*<0.0001. E. Western blot on whole-cell lysate from Sin3B^+/+^, Sin3B^-/-^ and Lin9^-/-^ T98G cells expressing different Sin3B truncation mutants probed using the indicated antibodies. F. Sin3B^+/+^, Sin3B^-/-^ and Lin9^-/-^ T98G cells were treated with 25 ug/mL of cisplatin for 12 hours. Cells were then subjected to Annexin V staining followed by flow cytometry analysis. Annexin V positive cells were then quantified. Error bars represent SD; n = 3. Significance was determined by two-way ANOVA followed by a Tukey’s test. \*\*\*\**P*<0.0001.

**Figure S3.**
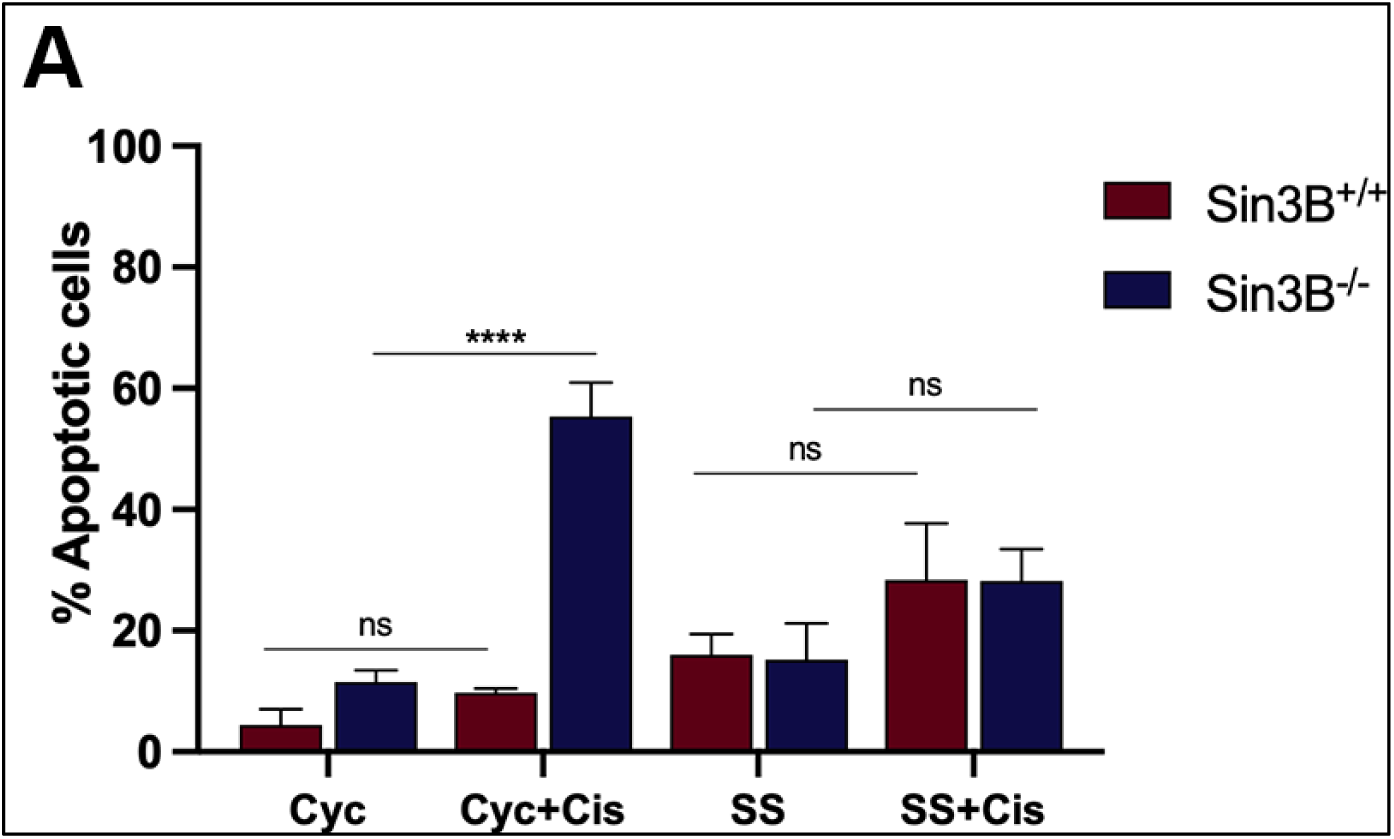
Cell cycle exit protects against DNA damage-induced apoptosis. A. Sin3B^+/+^ and Sin3B^-/-^ T98G cells were synchronized in G_0_ phase of the cell cycle by serum starvation for 72 hours before being treated with 25 ug/mL of cisplatin for 20 or 40 hours. Cells were then subjected to Annexin V staining followed by flow cytometry analysis. Annexin V positive cells were then quantified. Error bars represent SD; n = 3. Significance was determined by two-way ANOVA followed by a Tukey’s test. \*\*\*\**P*<0.0001.

**Figure S4.**
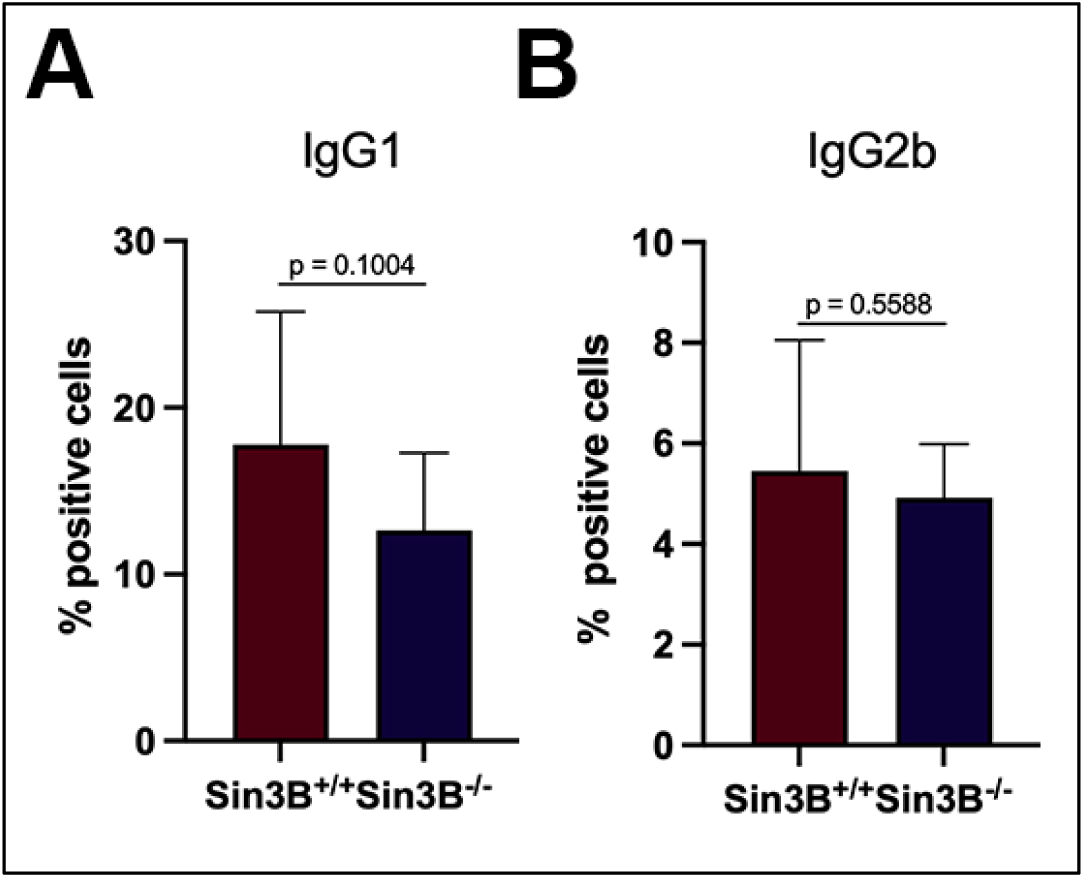
Sin3B^-/-^ mouse B cells exhibit mild defect in Class Switch Recombination. A. Splenic B cells from Sin3B^-/-^ mice were isolated and stimulated to undergo class switching with LPS (20 ug/ml) and IL-4 (10 ng/ml). Surface IgG1 expression was assessed on day 4 by flow cytometry. Error bars represent SD; n = 9. Significance was determined by Student’s t-test. B. Splenic B cells from Sin3B^-/-^ mice were isolated and stimulated to undergo class switching with LPS (20 ug/ml). Surface IgG2b expression was assessed on day 4 by flow cytometry. Error bars represent SD; n = 9. Significance was determined by Student’s t-test.

**Figure S5.**
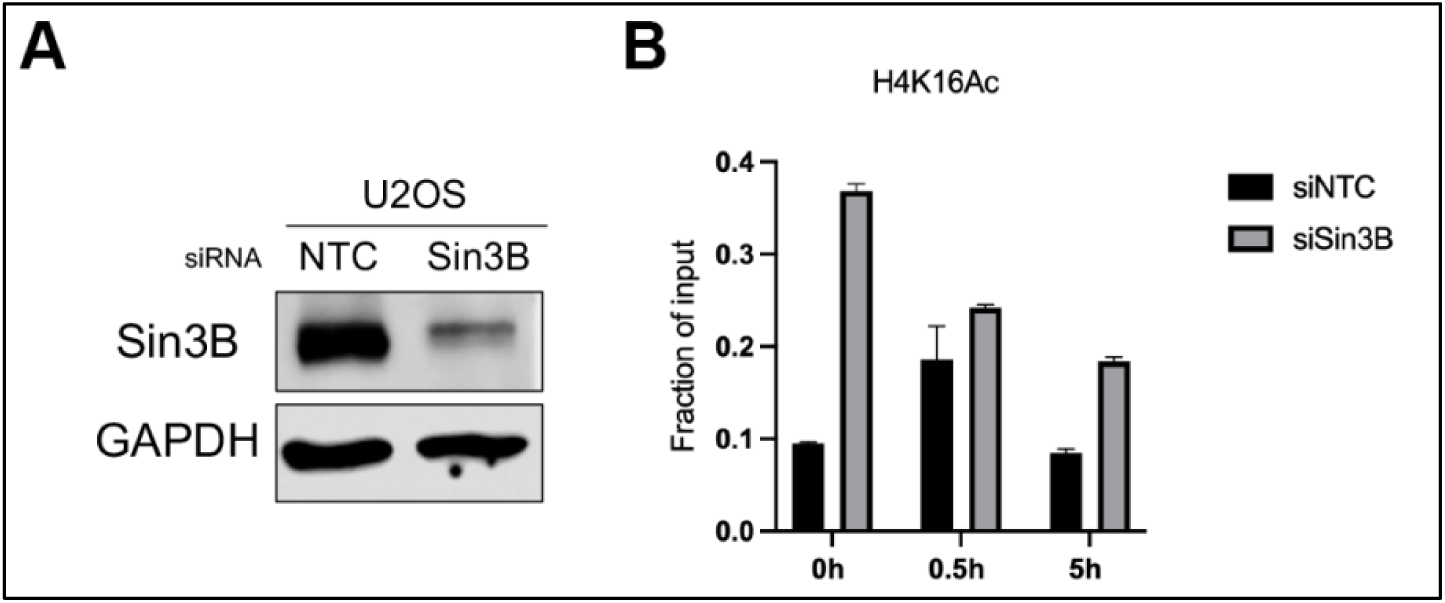
Sin3B deletion results in unchanged H4K16 Acetylation levels upon DSB induction. A. Western blot on whole-cell lysate from U2OS cells harvested 72 hours after siRNA transfection, probed using the indicated antibodies. B. ChIP-qPCR performed with antibody to H4K16Ac in the U2OS-DSB reporter cells. DSBs were induced 72 hours after siRNA transfection. Error bars represent SD; n = 3.

